# The tissue-specific chromatin accessibility landscape of *Papaver somniferum*

**DOI:** 10.1101/2022.04.13.487984

**Authors:** Yu Xu, Yanyan Jia, Bo Wang, Li Guo, Mengyao Guo, Xiaofei Che, Kai Ye

## Abstract

Accessible chromatin regions (ACRs) at promoters, enhancers, and other gene regulatory regions allow transcription factors (TFs) to bind, which regulate gene transcription involved in plant development and metabolism. *Papaver somniferum* has been widely applied in clinical medicine as one of the most important and oldest medicinal plants due to its unique and effective active ingredients. However, the transcriptional regulatory mechanism of tissue-specific distribution of active ingredients remains unknown. In this study, transcriptome and chromatin accessibility analysis by RNA sequencing (RNA-seq) and assay for transposase-accessible chromatin sequencing (ATAC-seq) was performed to investigate these underlying molecular mechanisms. We identified tissue-specific chromatin Tn5 hypersensitive site (THS) and gene expression by examining the variation of THS and transcripts across six tissues (capsule, stem, fine root, tap root, leaf, and petal). Our results provide insight into the epigenetic mechanism of transcriptional plasticity for *P. somniferum* organ development. Sequence motif analysis within accessible chromatin regions for co-expressed gene modules revealed enriched binding sites of hub transcription factors that regulate tissue-specific functions. Furthermore, we identified regulatory elements for tissue-specific accumulation of morphine and noscapine in *P. somniferum*. This is the first tissue-specific chromatin accessibility landscape of *P. somniferum* providing an important resource for functional epigenetic analysis and future molecular breeding in *P. somniferum* for variety improvement.

## INTRODUCTION

Opium poppy (*Papaver somniferum*) is a flowering plant in the Papaveraceae family that has been valued for its ornamental and significant medicinal properties for thousands of years (Norn *et al*., 2005, Singh *et al*., 2019). It produces several pharmacologically active benzylisoquinoline alkaloids (BIAs), such as morphine, codeine, thebaine, and noscapine, which make poppy the only natural source of commercial opiates worldwide and potentially play roles in plant defense against biotic and abiotic challenges. Developmental regulation of BIA biosynthesis facilitates organ and tissue-specific accumulation of major alkaloids. The primary alkaloids are mainly accumulated in the stems and capsules of mature plants (Facchini and De Luca, 1995, Hagel and Facchini, 2013, Beaudoin and Facchini, 2014). However, the regulatory mechanisms behind the tissue-specific production and enrichment of natural products in opium poppy are largely unknown. In addition, tissue-specific control of physiology, development, and growth of opium poppy is also unknown. Previously our research have reported that most genes that encode enzymes for metabolic pathways of BIAs are not only clustered on the poppy genome but also co-expressed in stem, capsule and root tissues (Guo *et al*., 2018, Yang *et al*., 2021). The mechanism by which co-expression of BIA genes occurs selectively in some tissues but not in others is intriguing and unknown. Therefore, we aimed to sequence and study the epigenome and transcriptome of distinct opium poppy tissues to uncover the tissue-specific regulating mechanisms of general plant physiology and, in particular, BIA production,

Accessible chromatins regions (ACRs) located at promoters, enhancers, and other gene regulatory regions allow transcription factors (TFs) to bind, which is crucial for transcriptional regulation during a wide range of developmental and metabolic processes (Thurman *et al*., 2012, Yocca and Edger, 2022). At present, the assay for transposase accessible chromatin sequencing (ATAC-seq) is being developed as an alternative approach for detecting the highly ACRs and then identifying TF-binding sites within these regions. ATAC-seq has been widely employed in recent years for large-scale identification of open chromatin in mammals, fungi and plants as a quicker and more efficient approach (Lu *et al*., 2017, Maher *et al*., 2018, Sijacic *et al*., 2018, Klemm *et al*., 2019, Pawlak *et al*., 2019, Chen *et al*., 2021), however, there has been no report regarding *P. somniferum*. Herein, we conducted comprehensive tissue-specific assay for ATAC-seq and transcriptome sequencing (RNA-seq) analysis of six different tissues in opium poppy to dissect the epigenetic and transcriptional regulation mechanisms for plant development and secondary metabolism. In this study, we not only identified tissue-specific ACRs, gene expression, and hub transcription factors in the scope of the poppy genome but also discovered that HB6 was a key transcription factor to regulate the expression of the BIA gene cluster. This first tissue-specific chromatin accessibility landscape of *P. somniferum* provides an important resource for functional epigenetic analysis and future research aimed at characterizing or using gene regulatory elements for breeding poppy varieties with high content of BIAs.

## RESULTS

### Tissue-specific transcriptomics of *P. somniferum* reveal transcriptional plasticity

To understand the tissue-specific transcriptional regulation in opium poppy, we performed RNA-seq of six different tissues of *P. somniferum* HN1 variety, including the leaves, petals, stems, capsules, tap roots, and fine roots. The tissues were all harvested on the first day of anthesis. After quality control, the RNA-seq data were aligned to the reference genome of *P. somniferum* (Yang *et al*., 2021), followed by transcript discovery, quantification, and normalization (see Materials and Methods). Overall, we found that RNA-seq data had high mapping ratios (between 75% and 80%) against the reference genome of *P. somniferum* (**Supplementary Table 1**). Gene expression was highly correlated in three biological replicates per tissue, with an average Pearson correlation coefficient (PCC) > 0.857 (**Supplemental Figure 1**), showing the high quality of the RNA-seq dataset. Based on gene expression profiles, the principal component analysis (PCA) showed a clean separation of various tissue types (**Figure 1A**). Both correlation analysis (**Supplemental Fig. 1**) and PCA (**Figure 1A**) revealed a clear transcriptome reprogramming associated with distinctive *P. somniferum* tissues. The correlation pattern of gene expression among the tissues is overall consistent with the closeness of their spatial distribution. For example, the fine roots and tap roots, which were strongly correlated in gene expression, were located close to each other on the PCA plot but distant from the other tissues (**Figure 1B)**, whereas the petals showed the lowest correlation with any other tissue. With a threshold of transcripts per kilobase per million mapped reads (TPM) larger than 1, the number of expressed genes per tissue ranged from 24537 (petals) to 31768 (capsules), representing approximately 44% to 57% of the total genes of *P. somniferum*, respectively (**Supplementary Table 2**). Comparison of the expressed genes among the tissues shows that approximately 36% of the genes (19682) were expressed in all the six tissues **(Figure 1B)**, with each tissue having various numbers (0.5% to 1.2%) of uniquely expressed genes. The capsules contained the most tissue-specifically expressed genes (663), out of the six tissues studied (**Figure 1B).** The fact that the capsule has the most expressed and unique genes of any tissue underlines the transcriptional and physiological hyperactivity in this tissue.

**Figure 1.**
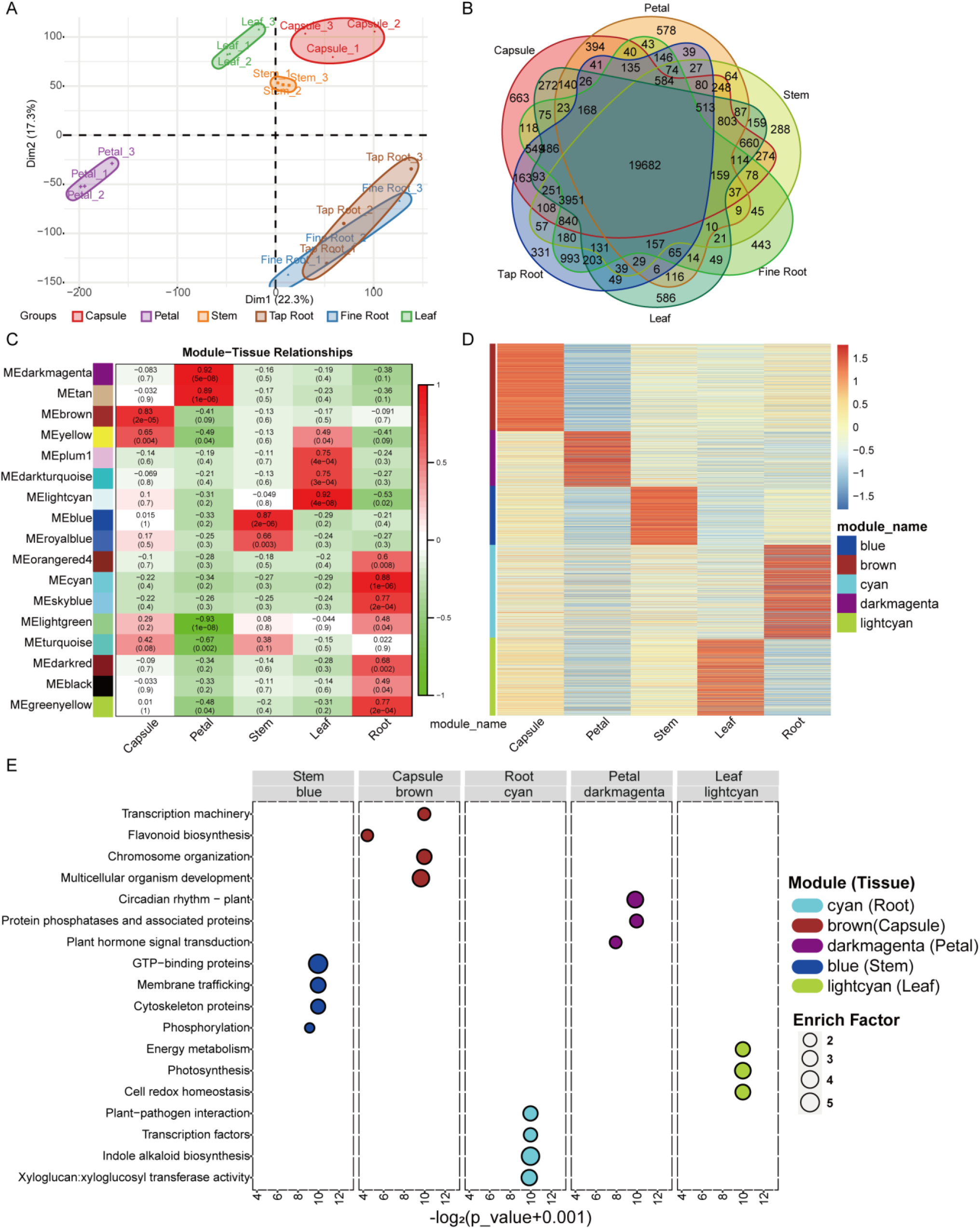
The transcriptome landscape of *P. somniferum* in six different tissues (capsule, leaf, petal, stem, tap root, and fine root). (A) Graphical representation of PCA for RNA-seq data of *P. somniferum*. (B) Venn diagram displaying overlaps between the expressed genes in the six different tissues. (C) Modules identified by WGCNA using RNA-seq data. (D) Heatmap of gene expression profiles in each tissue-specific functional module. (E) Enrichment analysis (GO and KEGG) of each tissue-specific functional module.

We performed a whole-genome co-expression network analysis (WGCNA) to identify the co-expressed gene modules responsible for tissue-specificity and differentially investigate the regulated genes underpinning the observed transcriptional plasticity (Langfelder and Horvath, 2008). The WGCNA revealed 17 co-expressed gene modules (CGMs) using expression data from the six tissues (**Supplementary Table 3**), each showing a significantly preferential association with certain plant tissues (*p* value < 0.05; correlation coefficient (*r*) > 0.5) (**Figure 1C**). Five modules were considered tissue-specifically associated gene modules (the smallest *p* value), *i.e*., the capsule module (brown, 4635 genes), petal module (dark-magenta, 2766 genes), stem module (blue, 2899 genes), leaf module (light-cyan, 3817 genes), and root module (cyan, 4688 genes) (**Figure 1C**). Expectedly, these five modules showed higher gene expression in their associated tissues than in the other tissues (**Figure 1D**). Functional enrichment analysis showed that the five modules were enriched in quite distinctive biological processes (**Figure 1E**). For instance, the capsule module was enriched in spliceosome, messenger RNA biogenesis, chromosome and associated proteins, flavonoid biosynthesis, and developmental process. The petal module was enriched with protein phosphatases and associated proteins, circadian rhythm, autophagy, and plant hormone signal transduction. The stem module was enriched with membrane trafficking, GTP-binding proteins, cytoskeleton proteins, and phosphorylation. Unsurprisingly, the leaf module was mainly enriched in photosynthesis. The root module was enriched in indole alkaloid biosynthesis, plant-pathogen interaction, and xyloglucosyl transferase activity **(Supplemental Table 4)**. We identified five gene modules with tissue-specific expression patterns and biological functions, reflecting the diverse transcriptional regulatory networks that underpin the distinctive tissues of *P. somniferum*.

### Identification of accessible chromatin regions in *P. somniferum* by ATAC-seq

We performed chromatin accessibility profiling using ATAC-seq from the same six tissues of which transcriptomes were assessed (see Materials and Methods). The purpose of this analysis was to map genome-wide cis-regulatory elements involved in transcriptional regulation in *P. somniferum*. The ATAC-seq libraries of three biological replicates for each tissue were sequenced using Illumina paired-end sequencing, yielding a total of 890 million clean reads that were mapped to the reference genome of *P. somniferum* variety HN1 (**Supplementary Table 5**). Similar to the PCA of the transcriptomic profiles, the PCA of the ATAC-seq data showed that the three biological replicates within each tissue were highly correlated and roughly separated into tissue-specific clusters (**Figure 2A**), with the exception that the clusters of tap root and fine root intermingled with each other (**Figure 2A**), reflecting a correlation of the two tissues in both chromatin accessibility and gene transcription.

**Figure 2.**
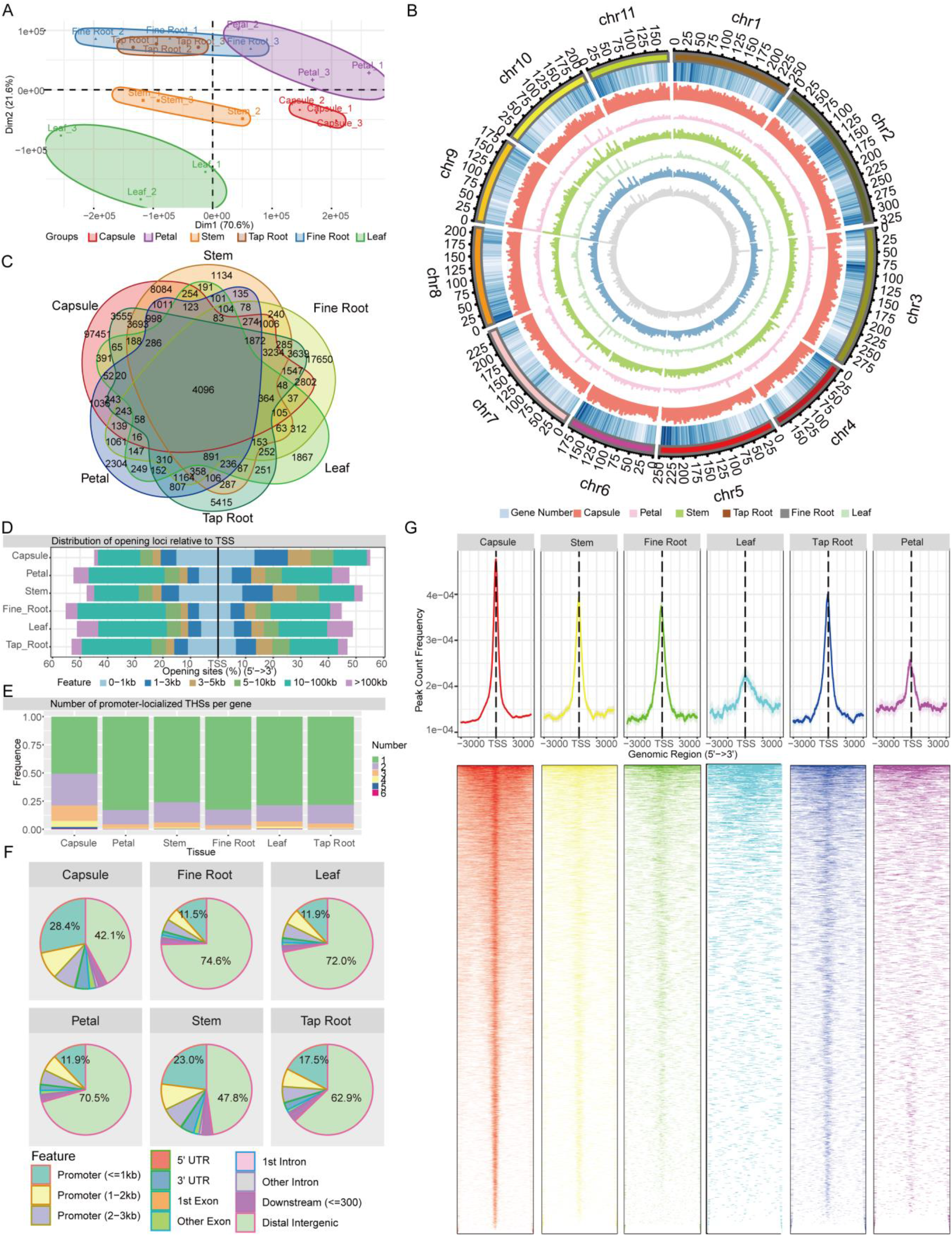
Landscape of accessible chromatin regions across six tissues (capsule, leaf, petal, stem, tap root, and fine root) of *P. somniferum*. (**A**) Graphical representation of PCA of opium poppy ATAC-seq data between all the biological replicates across the six tissues based on THS. (**B**) Genome-wide distribution of THSs along 11 poppy chromosomes. From outside to inside, the circles represent the chromosome, gene density, THSs abundance in the capsule, THSs abundance in the petal, THSs abundance in the stem, THSs abundance in the leaf, THSs abundance in the tap root, and THSs abundance in the fine root. (**C**) Venn diagram displaying the number of common and unique THSs among the six distinctive tissues. (**D**) Distribution of THSs relative to genes in each tissue. (**E**) Number of promoter-localized THSs per gene in each tissue. (**F**) Genomic annotation and distribution of THSs in the six tissues. (**G**) Distribution of promoter-localized THSs in the six tissues.

For each tissue, we identified statistically significant ATAC-seq peaks (*p value* < 0.01) representing ACRs specifically enriched with THS using Genrich software (https://github.com/jsh58/Genrich). As a result, 133374, 18914, 31974, 11525, 35877, and 43565 THSs were identified in the capsules, petals, stems, leaves, tap roots, and, fine roots, respectively (**Figure 2B and 2C; Supplemental Table 6**). The presence of most of THSs and expressed genes in the capsules (**Figure 1B**) indicated that transcriptomic changes in the capsule are positively correlated with open chromatin states. The comparison of THSs in the different tissues showed that 4096 THSs were shared by all the tissues, with the capsule having the most tissue-specific THSs (**Figure 2C**). Overall, THS distribution along *P. somniferum* chromosomes resembles the gene density distribution (**Figure 2B**), showing that THSs are more frequently located near gene-dense regions rather than near gene-sparse regions. Examining the location of THSs relative to genes showed that in all the tissues, except the capsule (46.1%) and stem (40.3%), approximately 20% to 30% THSs were located within 3kb regions of the transcription start site (TSS) (**Figure 2D**), while 42.1% to 74.6% of THSs were located in distant intergenic regions. In comparison to other tissues, the capsule (28.4%) and stem (23.0%) contained more THSs within 1kb promoter regions and fewer THSs in distal intergenic regions (**Figure 2F**). As for the genes with a detected promoter region of the THSs across the six tissues, 80%, 12% to18%, and 3% to 5% had a single THS, two THSs, and three THSs, respectively, except for the capsule, which had 50%, 28%, and 14%, respectively (**Figure 2E**). The peak of the promoter-localized THS was located around TSS for all the six tissues, with 61.47%, 56.95%, 58.87%, 55.8%, 52%, and 51.74% within the 1kb of TSS in the capsule, stem, tap root, fine root, leaf, and petal, respectively, demonstrating a more open chromatin state around TSS than the rest of promoter regions (**Figure 2G**). Taken together, the ATAC-seq detected a large amount of THSs with a distinct distribution associated with six tissues, reflecting a common and distinct state of open chromatins among these tissues.

### Chromatin accessibility is correlated with gene expression pattern

We next investigated the correlation of detected ACRs with gene expression profiles across different tissues. Comparing promoter-localized THSs (pTHSs) of six tissues revealed 59920, 2959, 11043, 1244, and 4009 tissue-specific pTHSs (defined as pTHSs per tissue minus pTHSs shared by all tissues) were identified in capsule, petal, stem, leaf and root, respectively (**Figure 3A**). Overall, genes associated with these tissue-specific pTHSs showed significantly higher (average log2FC > 2.85 and *p* value < 2.22e-16) expression than those without (**Supplemental Figure 2**). Such pTHS-associated genes were significantly (*p* value < 0.05, Fisher’s exact test) enriched in tissue-specific WGCNA modules (**Figure 3B; Supplemental Figure 3**) for all tissues examined. This implies an association between the identified ACRs and expression profiles in specific tissues with a remarkable functional relevance. For example, a *P. somniferum* ortholog (*Pso06G14270.0*) of *Arabidopsis STM*(Homeobox protein SHOOT MERISTEMLESS), participated in shoot apical meristems maintenance (Aida *et al*., 1999), was highly expressed in the stem belonging to the blue (stem) module, possessing a stem-specific pTHS that is absent in the other tissues (**Figure 3C**). Likewise, the *YAB3* (axis regulator YABBY 3, *Pso07G00980.0*), involved in fruit development (Dinneny *et al*., 2005), *LHCB1.2* (chlorophyll a-b binding protein 3, *Pso06G37060.0*), in photosynthesis (Sun and Tobin, 1990), *COP1* (E3 ubiquitin-protein ligase COP1, *Pso06G14590.0*), in photomorphogenesis and circadian rhythm (Lau and Deng, 2012), and *MAPKKK20 (Pso01G36990.0*), in primary root cell division elongation (Benhamman *et al*., 2017, Li *et al*., 2017), showed preferential expression in the capsule, petal, leaf, and root, respectively and carried the specific pTHSs in the corresponding tissues (**Figure 3C**). Moreover, we identified 803 pTHSs shared by the six tissues (**Figure 3A**) that were localized in the promoter regions of 664 genes, 340 of which were expressed (average TPM > 1). Functional enrichment showed that the 340 genes are prominently enriched in fundamental biological processes such as plant hormone signal transduction, oxidative phosphorylation, messenger RNA biogenesis, and energy metabolism (**Supplemental Table 7**). For instance, *RGA* (DELLA protein RGA, *Pso05G17050.0*) plays a negative role in gibberellin (GA) signaling pathway (Silverstone *et al*., 2001), whereas *PFK5* (ATP-dependent 6-phosphofructokinase 5, *Pso04G23600.0*) participates in the first committed step of glycolysis (Mustroph *et al*., 2007). These genes carried common pTHSs and were expressed comparably in the six tissues (**Figure 3D**). Ultimately, co-profiling of chromatin accessibility and transcriptome displays a strong correlation between ACRs and associated gene expression, indicating that differences in accessible chromatin states could well explain the tissue-specific transcriptomic and functional plasticity observed in *P. somniferum*.

**Figure 3.**
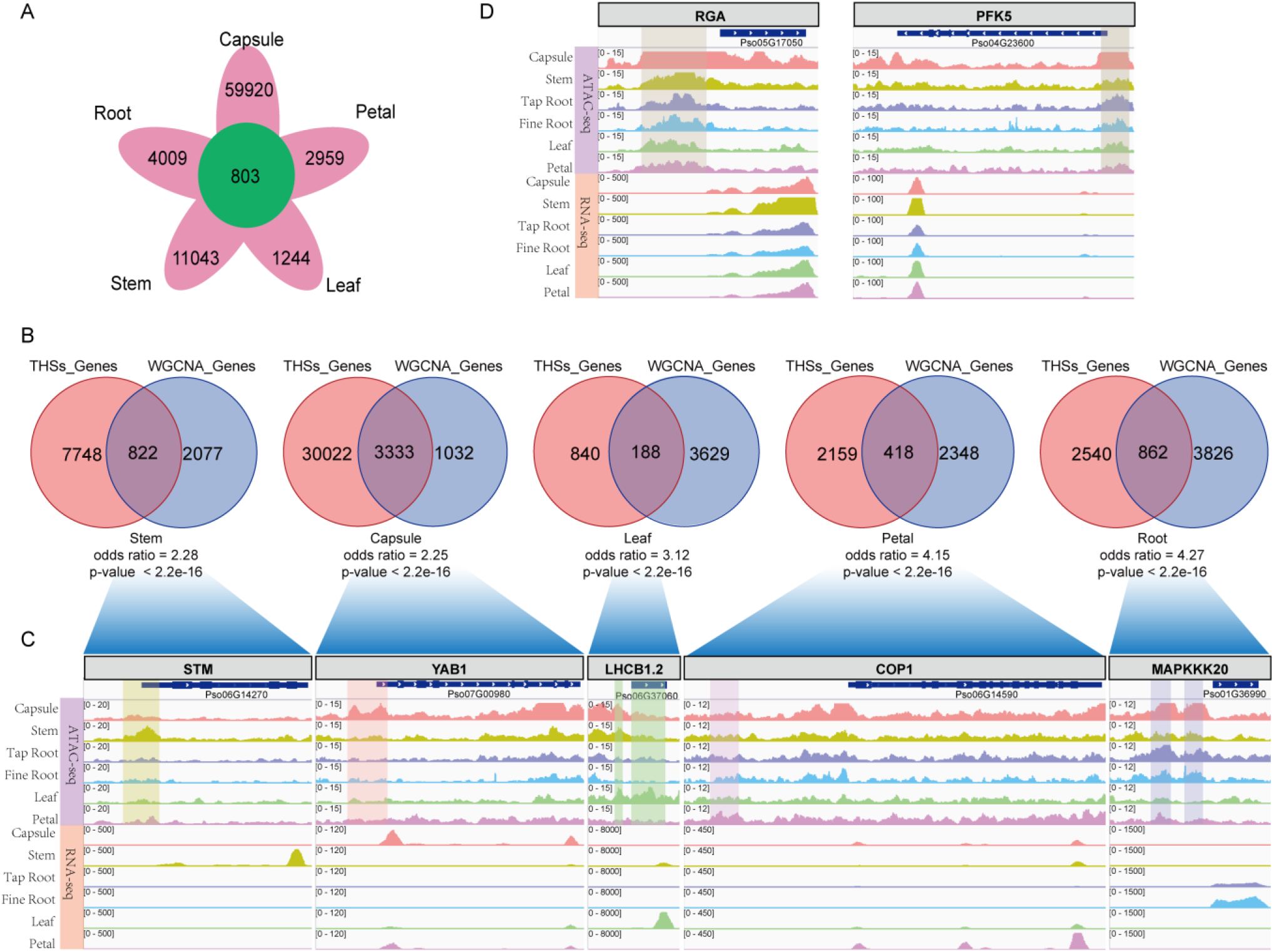
Co-identification of open chromatin regions and gene expression with tissue-specificity in *P. somniferum*. A. Circle diagram showing the number of THS shared (green circle) and specific (magenta cycles) to each tissue. B. Venn diagrams, each presenting a comparison between genes close to tissue-specific THS and genes within tissue-specifically co-expressed gene modules. The odds ratio and *p*-value represent the enrichment factor and statistical test for each tissue. C. Genes with both tissue-specific expression and proximity to a tissue-specific THS are visualized in IGV, with the THS peak and RNA-seq peak identified at or around the gene coding sequence.

### Joint analysis of RNA-seq and ATAC-seq identifies tissue-specific hub transcription factors

ACRs provide TFs with access to cis-regulatory elements, enabling fine-tuned transcriptional regulation through TF-DNA binding. With the identified ACRs, we sought to characterize potential cis-regulatory elements and associated TFs through a computational pipeline that integrates sequence motif enrichment in pTHSs, TF binding motif (TFBS) similarity assessment, and co-expression network analysis (see Methods). The logic is that TFs that bind to TFBS are located in ACRs of target genes with which they are often co-expressed. First, *P. somniferum* genes associated with both tissue-specific pTHSs and a WGCNA module (**Figure 3B**) were used to identify significantly enriched DNA motifs using HOMER software (Heinz *et al*., 2010) based on the tissue-specific pTHS sequence. Second, highly expressed (TPM > 5) TFs within the WGCNA module (**Figure 1C**) were considered as candidate hub TFs for the module. Lastly, the candidate hub TFs were considered highly confident when motifs matched between enriched motifs of tissue-specific pTHS and reported motifs associated with the candidate hub TFs.

WGCNA module genes carrying tissue-specific pTHSs were highly expressed in their representative tissue, as illustrated in **Figure 4A**. Using the aforementioned developed computational pipeline on such genes, we identified seven, seven, fourteen, eight, and one hub-TF in *P. somniferum* co-expression modules for the capsule, stem, root, petal, and leaf, respectively (**Figure 4B; Supplemental Table 8**). These hub-TFs putatively bind to their TFBS, which are found in ACRs associated with target genes in corresponding tissues.

**Figure 4.**
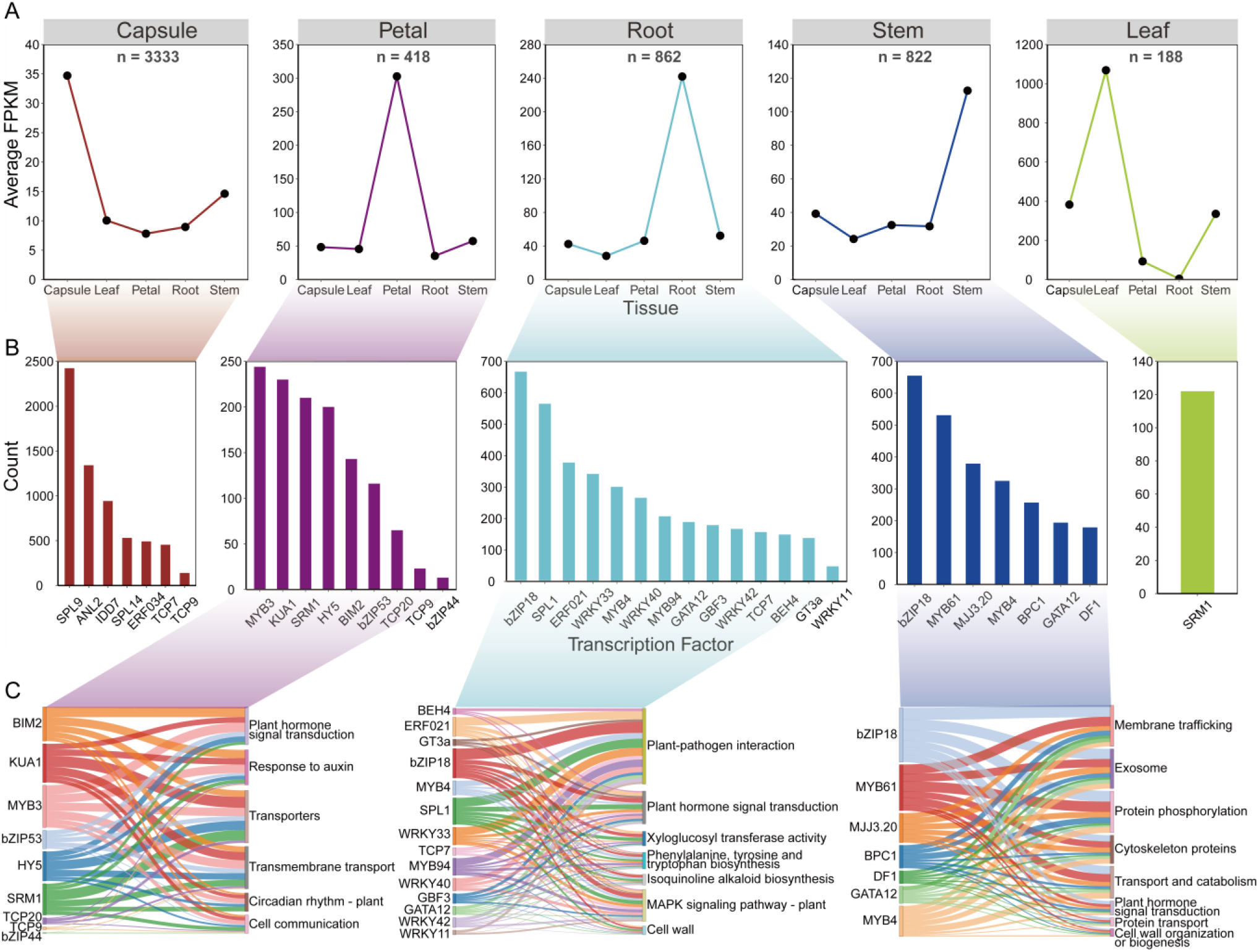
THSs identifies regulatory elements and dynamic regulatory networks of different tissues. (A) Expression profiles of the 5623 genes identified in Fig 3B. (B) Number of genes regulated by each hub transcription factor in five tissues. (C) Regulatory relationships between transcription factors and enrichment terms in the petal, root, and stem tissues.

Seven hub-TFs have been identified for the capsule module, including *SPL9, SPL14, ANL2, ERF034, IDD7, TCP7*, and *TCP9*, which together regulate 2740 genes enriched with functional terms, such as chromatin organization and modification, developmental processes, and diverse metabolic functions (*e.g*., flavonoid, carbon, lipid, amino acid, vitamins) (**Supplemental Table 9 and 10**). Among these seven hub-TFs, *SPL9* was found to be the most influential hub-TF regulating 2423 (88.4% of 2740) genes (**Figure 4B**), including genes involved in flavonoid biosynthesis, such as chalcone isomerases, chalcone synthases, shikimate O-hydroxycinnamoyltransferases, and leucoanthocyanidin reductases. In addition, we found that *SPL9* also regulated a *P. somniferum* bifunctional dihydroflavonol 4-reductase/flavanone 4-reductases (DFR), which is consistent with reports on annotated *DFR* promoter binding-affinity function of *SPL9* (Gou *et al*., 2011b). Moreover, *SPL9* regulated thirteen pectinesterase genes, eight of which were preferentially expressed in the capsule (average TPM = 111.5), suggesting their functional importance for fruit development (**Supplemental Figure 4**).

For the petal module, eight hub-TFs (*i.e., SRM1, MYB3, KUA1, HY5, BIM2, bZIP44, bZIP53, TCP9*, and *TCP20*) were found; these transcription factors regulated 382 genes enriched in plant hormone signal transduction, circadian rhythm, and transmembrane transport (**Figure 4C; Supplemental Table 9 and 10**). As shown in **Figure 4B**, the four most enriched hub-TFs were *MYB3, KUA1, SRM1*, and *HY5*, each of which regulated at least 200 genes associated with key petal functions (**Figure 4B; Figure 4C**). Circadian rhythm is a significant function for petal tissue. Using our exhaustive transcriptional regulatory network, we found *MYB3* and *SRM1* together with some other TFs to regulate these circadian rhythm-related genes, highlighting the utility of our transcriptional regulatory network (**Figure 4C; Supplemental Table 11**).

Using root-specific promoter-localized THSs shared by tap root and fine root, we identified fourteen hub-TFs in the root module, which regulated 837 genes with functions related to plant-pathogen interaction, plant hormone signal transduction, xyloglucosyl transferase activity, cell wall, and phenylalanine, tyrosine, and tryptophan biosynthesis (**Figure 4B and 4C; Supplemental Table 9 and 10**). *bZIP18* was the most enriched of these fourteen hub-TFs, regulating over 650 genes involved in plant-pathogen interaction, plant hormone signal transduction, xyloglucosyl transferase activity, cell wall, phenylalanine biosynthesis, isoquinoline alkaloid biosynthesis, *etc*. Interestingly, bZIP18, as one of seven stem hub-TFs, had the highest number of regulated genes involved in transport and catabolism, membrane trafficking, exosome, plant hormone signal transduction, cell wall organization, *etc*. (**Figure 4B and 4C**; **Supplemental Table 12**). These results implied that although *bZIP18* played an important regulatory role in the roots and stems, the focus of regulation is different.

*SRM1* was the only hub-TF in the leaf module that regulated 122 genes, 39 of which were photosynthesis-related proteins (**Figure 4B; Supplemental Table 9 and 10**). Nevertheless, the core function of *SRM1* in the petal was to regulate circadian rhythm-related genes (**Supplemental Table 11 and 13**), including *COP1, SAP1*, and *HY5*. As a result of gene duplication and subsequent functional divergence, our findings reveal that different paralogs of the common transcription factor regulate distinctive functional genes in different tissues.

In summary, the diverse regulatory networks of these various transcription factors in different tissues result in distinctive tissue-specific functions.

### Tissue-specific chromatin accessibility and transcription of BIA gene cluster

*P. somniferum* production of pharmaceutically valuable BIA, such as morphine, codeine, thebaine, and noscapine is a distinguishing characteristic (Beaudoin and Facchini, 2014). The genes encoding the BIA biosynthetic pathway are well characterized and organized in a gene cluster on the *P. somniferum* genome (**Figure 5A**) (Guo *et al*., 2018, Yang *et al*., 2021). The fact that these BIA genes are transcriptionally co-regulated in a tissue-specific (capsule, root, and stem) manner (**Supplemental Figure 5**) raises the question of how this is achieved epigenetically and what regulatory elements are involved. Using enrichment analysis of DNA cis-regulatory elements in ATAC-seq data, we detected 43 capsule-, stem-, and root-specific pTHSs 3kb upstream of most BIA biosynthesis genes that encode the (*S*)-reticuline pathway (*NCS, 6OMT, CNMT, CYP80B1*, and *4OMT*) and sequentially convert L-dopamine and 4-HPA into the (*S*)-reticuline, morphinan branch (*STORR, Salsyn, SalR, SalAT, THS*, and *COR*), and noscapine branch (*PSMT1, CYP719A21, TNMT, CYP82Y1*, and *CYP82X2*) pathways, which is in accordance with their capsule-specific, root-specific and stem-specific gene expression (**Figure 5B; Supplemental Table 14**). In contrast, we did not observe any THSs of these genes in the non-BIA producing tissues, such as the leaf and petal, where they are lowly expressed (**Figure 5B**). These findings provide evidence that the chromatin becomes accessible to certain unknown transcriptional regulators (*e.g*., TFs) in specific tissues allowing them to activate the transcription of many of these BIA genes.

**Figure 5.**
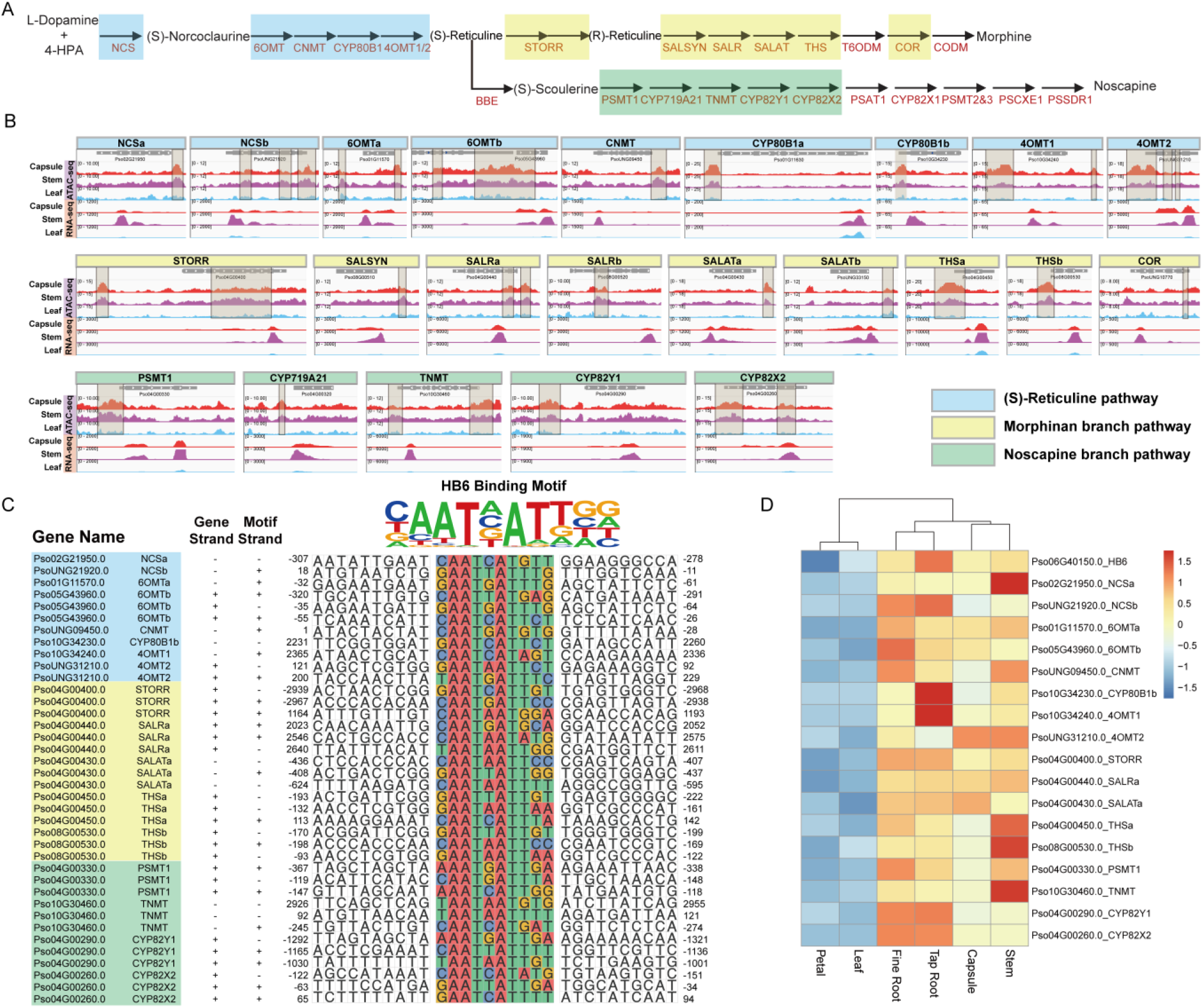
Coordinate regulation of BIAs metabolic pathway genes. (A) Schematic representation of (*S*)-reticuline pathway and noscapine and morphinan branch pathways. (B) Visualization of THSs from −3000 to 3000 bp relative to the TSS of these isoquinoline metabolic pathway genes. (C) The predicted cis-regulatory element within the region from −3000 to 3000 bp relative to the TSS of (*S*)-reticuline, noscapine, and morphinan metabolic pathway genes. (D) The expression patterns of HB6 as well as its predicted regulated genes.

Subsequently, we identified potential TFBS from the 43 pTHSs associated with BIA genes through motif enrichment analysis (**Figure 5B; Supplemental Figure 6 and Table 14**). The HB-HD-ZIP family and the WRKY family recognition motifs were significantly enriched among these pTHSs, which is consistent with previous studies on the function of WRKY family proteins in the regulation of BIA biosynthesis in California poppy (*Eschscholzia californica*) (Yamada *et al*., 2021). Nevertheless, combining the results of motif enrichment and RNA-seq analysis, HB6, a HB-HD-ZIP transcription factor, emerged as a key regulator in BIA biosynthesis in *P. somniferum*. Specifically, *Pso06G40150.0*, which encodes a *P. somniferum* homolog of *Arabidopsis* HB6, regulated 19 BIA biosynthetic genes including eight in the (*S*)-reticuline pathway, five in the morphinan branch pathway, four in the noscapine branch pathway, one (P6H) in the sanguinarine branch pathway, and one (7OMT) in the laudanine branch pathway (**Figure 5C and Supplemental Figure 7**). HB6 was also highly co-expressed with these 19 genes (average correlation coefficient > 0.751) (**Figure 5D and Supplemental Table 15**). For instance, *Pso04G00400.0 (STORR*) is a pivotal gene in the morphine branch pathway (Winzer *et al*., 2015), which contained HB6 binding motif within its promoter region. This motif was significantly opened in the capsule, stem, fine root, and tap root, but nearly closed in the petal and leaf, which was consistent with its lower expression level in the petal and leaf (**Figure 5B and 5C; Supplemental Figure 8**). The remaining 18 genes yielded similar findings (**Figure 5B and 5C and Supplemental Figure 7**). In summary, these results suggested that *Pso06G40150.0* was a key transcription factor in regulating the expression of the BIA gene cluster.

Taken together, our findings suggest that several major genes involved in the BIAs metabolic pathway may be regulated coordinately by the same transcription factor.

## DISCUSSION

*P. somniferum*, one of the most important medicinal plants in the world, has been widely used in clinical medicine for thousands of years due to its unique ability to produce a variety of active ingredients, including noscapine, morphine, and codeine, all of which have potential pharmacological activity in relieving pain, cough, muscle relaxation, anticancer, *etc*. (Yamada *et al*., 2021). However, the regulatory mechanisms governing its development and tissue-specific product synthesis remain unclear. These research status limit full utilization and breeding improvement of *P. somniferum*.

Eukaryotic gene expression is typically regulated by the binding of various TFs to *cis*-elements located in ACRs, such as promoters, enhancers, and other gene regulatory regions (Thurman *et al*., 2012, Yocca and Edger, 2022). Mapping the *cis*-regulatory elements in ACRs is therefore crucial to understanding the landscape of dynamic gene regulation that underlies plant development and metabolism. ATAC-seq method, which uses a hyperactive Tn5 transposase that is preloaded with sequencing adapters as a probe for chromatin accessibility, provides a valuable technique for investigating the chromatin accessibility in the scope of the genome (Buenrostro *et al*., 2015). ATAC-seq has been successfully applied in mammals, certain plants, and fungi (Lu *et al*., 2017, Maher *et al*., 2018, Sijacic *et al*., 2018, Klemm *et al*., 2019, Pawlak *et al*., 2019, Chen *et al*., 2021).

In this study, we performed a combined ATAC-seq and RNA-seq analysis on six different tissues (*i.e*., capsule, stem, fine root, tap root, leaf, and petal) of *P. somniferum* separately to understand in detail how the synthesis of tissue-specific active ingredients is regulated and how the tissue specific-development is established at the regulatory level. The PCA for both RNA-seq and ATAC-seq, data as well as functional annotation for the tissue modules showed patterns associated with distinctive *P. somniferum* tissues. Since the tap root and fine root are extremely close to each other, we merged them into the same root module.

In our poppy ATAC-seq data, major ACRs were located in distant intergenic regions, which were significantly greater than in *Arabidopsis* (Maher *et al*., 2018, Sijacic *et al*., 2018). This difference may be caused by the highly compact genome of *Arabidopsis* and is consistent with the observation of a negative correlation between genome size and the distance of ACRs from genes reported by Maher (Maher *et al*., 2018). ACRs also vary between different tissues or cell types. For example, in a recent study of maize cis-element atlas, ACRs identified in the most enriched cell were 4.2-fold higher than average levels for all cells (Marand *et al*., 2021). In our study, there were dynamic changes in the ACR profile for six different tissues, notably for the capsule, which had the highest number of ACRs and specifically expressed genes. This outcome may be explained by high levels of metabolism and developmental activity. Our results are in agreement with those of prior research on tomato fruit (Qiu *et al*., 2016). Interestingly, we identified a large amount of covariation between ACRs and transcriptome profiles as well as a great number of specific transcription factors located in particular tissue-specific ACRs that may contribute to tissue specificity.

It has been reported that TFs bind to TFBS found in ACRs of target genes with which they are often co-expressed. Therefore, we performed a joint RNA-seq and ATAC-seq analysis to identify tissue-specific hub TFs. As the most influential hub-TF in the capsule module, *P. somniferum* homolog (*Pso09G17650.0* and *Pso11G22980.0*) of *SPL9* has been reported to encode a *SQUAMOSA PROMOTER BINDING PROTEIN-LIKE* protein in *Arabidopsis* to regulate various processes including anthocyanin and flavonoid production (Gou *et al*., 2011a). The same homolog presented in our study as the most influential hub-TF in the capsule module and regulated 305 genes likely involved in various metabolic pathways in the capsule. Moreover, eight capsule-preferentially expressed pectinesterase genes (average TPM = 111.5) were regulated by *SPL9*, which may play roles in fruit ripening through participating in cell wall metabolism (Phan *et al*., 2007). We noticed that *MYB3, KUA1*, *SRM1*, and *HY5* were the four highest enriched hub-TFs in the petal module, with *MYB3* and *SRM1* working together with some other TFs to regulate these circadian rhythm-related genes. These findings support the idea that circadian rhythm is a significant function for petal tissue. We also identified 14 hub-TFs in the root module using root-specific promoter-localized THSs shared by the tap root and fine root. These hub-TFs regulated a total of 837 genes with functions related to plant-pathogen interaction, plant hormone signal transduction, xyloglucosyl transferase activity, cell wall, and phenylalanine, tyrosine, and tryptophan biosynthesis (**Figure 4B and 4C**). We found two gene family clusters among these 837 genes, namely the jasmonate ZIM domain-containing (JAZ) gene family cluster (de Torres Zabala *et al*., 2016, Feng *et al*., 2020, Han *et al*., 2020) and xyloglucan endotransglucosylase/hydrolase (XTH) gene family cluster (Li *et al*., 2019, De Caroli *et al*., 2021) (**Supplemental Table 16**). Both the *JAZ* and *XTH* genes were extremely expressed in the roots (**Supplemental Figure 9**). One gene (*Ps02G24140.0*) of the *JAZ* gene family and one gene (*Ps06G25510.0*) of the *XTH* gene family were highly expressed and co-regulated by *SPL1, bZIP18*, and *ERF021* in the roots (**Supplemental Figure 10**). This outcome is also consistent with *SPL1, bZIP18*, and *ERF021* being the three transcription factors with the highest number of regulated genes in the roots (**Figure 4B**).

In addition to *XTH* genes being annotated as regulators of cell wall modification in the root, certain target genes of *bZIP18* in the stem were also associated with the cell wall, such as cellulose biosynthetic process (3 genes, *p-value* = 0.073) and cell wall organization or biogenesis (5 genes, *p-value* = 0.099) (**Supplemental Table 12**). The functional annotations of these proteins are related to plant growth and development and belong to secreted or membrane proteins (**Supplemental Table 17**). Therefore, we explored the stem module in greater depth to better understand the differences between root and stem tissues performing the cell wall biogenesis function. In the stem module, there were seven hub-TFs regulating 777 genes enriched in cytoskeleton proteins, membrane trafficking, exosome, plant hormone signal transduction, and various substances transport **(Figure 4B and 4C**). Significantly, *bZIP18* and *MYB61* showed the highest number of regulated genes in the stem, coinciding with the previously annotated xylem-formation function of *MYB61* (Romano *et al*., 2012) (**Supplemental Table 18**). In addition, many genes in various plant hormone signal transduction pathways related to cell enlargement, elongation, and division were highly expressed in the stem (**Supplemental Table 18 and Figure 11**). For example, *TIR1, SAUR*, and *AUX/IAA* genes, which are closely related to auxin (Luo *et al*., 2018), were highly expressed in the stem and could almost be regulated by *bZIP18*. Generally, these results implied that although *bZIP18* regulates cell wall biosynthesis-related genes in root and stem tissues, the purposes of cell wall biosynthesis between root and stem tissues are different, with one focusing on the defense response (Li *et al*., 2019, De Caroli *et al*., 2021) and the other on growth and development (Luo *et al*., 2018). According to the above-mentioned results, we found that the majority of these hub-TFs regulate their targets in a tissue-specific manner, even though many hub-TFs were shared between different tissues.

Noscapine and morphine production are symbolic features of *P. somniferum*. A previous study showed that genes encoding the biosynthetic pathways of these metabolites were scattered in the ancestral various chromosomes for *P. somniferum* and that some of these were organized in a gene cluster and gained co-expressed profiles within the recent 7.2 Mya (Yang *et al*., 2021). Across many different co-expressed metabolic gene clusters, the co-expression patterns are constructed by coordinately regulation of specific regulatory proteins, such as cucurbitacin biosynthesis in cucurbits, which was regulated by three bHLH transcription factors (Shang *et al*., 2014, Qiu *et al*., 2016, Bharadwaj *et al*., 2021). In addition, we found coordinately chromatin accessibility variations with gene expression profiles for BIAs biosynthetic genes, suggesting that the above-mentioned rules may also play a role in *P. somniferum*. Therefore, we are eager to know how our focused natural product synthesis pathways are regulated to achieve these co-expressed patterns. In this study, we found that the promoters of some key genes involved in noscapine and morphine biosynthetic pathways harbored the HB-HD-ZIP family and the WRKY family recognition motifs. The involvement of WRKY proteins in regulating BIA biosynthesis has been elucidated in many Papaveraceae species, *e.g*., through wound-induced and MeJA related regulation (Yamada *et al*., 2021). However, the implication or role of HB-HD-ZIP family transcription factors involved in BIAs biosynthesis has yet to be characterized. Our results revealed and demonstrated that 19 BIA biosynthetic genes might be co-regulated by HB6 and that they were highly co-expressed with HB6. Although the molecular mechanism underlying the regulation of BIAs biosynthesis by these critical factors is still unclear, our study partly explained the reason for BIA gene co-expression patterns through co-regulation. The current research has also uncovered a valuable list of potential coordinate regulatory proteins participating in BIAs biosynthesis through transcriptomic and chromatin accessibility genomic evidence.

Hence, we constructed the first cis-regulatory elements landscapes from six distinctive tissues and provided paired transcriptomic data in *P. somniferum*. Our approach, which combines chromatin accessibility profiling with transcriptome profiling, is practicable and precise for identifying cis-regulatory elements and building regulatory networks construction. Our data atlas will provide a valuable resource for the study of epigenetic mechanisms underlying plant development and secondary metabolism.

## METHODS AND MATERIALS

### Plant materials and growth conditions

Opium poppy cultivar HN1 seeds were sowed in a soil mix containing potting mix, vermiculite, and sand at a 2:1:1 ratio. The seeds were incubated in plant growth chambers under a 16-hour light and 8-hour dark cycle at 22 °C and 60 % humidity. Six different tissues, including the leaves, stems (2-4 cm below the capsule), capsules, petals, tap roots, and fine roots of opium poppy plants were harvested one-day post-anthesis for ATAC-seq and RNA-seq experiments.

### Nuclei isolation and ATAC sequencing

A quantity of 1 to 3 g from the six different fleshy tissues was ground into powder in liquid nitrogen and then the nuclei were isolated as described previously. The isolated nuclei were immediately resuspended in the Tn5 transposase reaction mix. The transposition reaction was incubated at 37 °C for 30 min. Equimolar Adapter 1 and Adapter 2 were added after transposition and then PCR was performed to amplify the library. After PCR, the libraries were purified with the AMPure beads and library quality was assessed using Qubit. The library preparations were sequenced after cluster generation on an Illumina Hiseq platform and 150 bp paired-end reads were generated. The ATAC-seq was performed by Tianjin Novogene Technology Inc.

### RNA extraction and sequencing

The plant materials used for RNA-seq analysis were the same as those used for ATAC-seq. Total RNA was extracted from six different tissues using Trizol reagent (TaKaRa) according to the manufacturer’s instructions. RNA integrity was determined using regular agarose gel electrophoresis, Nanodrop (ThermoFisher Scientific), and Agilent 2100 Bioanalyzer (Agilent Technologies). RNA sample of high quality (OD260/280 within the range [1.8, 2.2], OD260/230 ≥ 2.0, RIN ≥ 8) was used to construct the sequencing library. A total amount of 1 μg RNA was used to build the library with Illumina Trueseq RNA Library Preparation Kit following the company’s recommended protocols. The RNA-seq libraries were sequenced on an Illumina Hiseq platform and 150 bp paired-end reads were generated at Tianjin Novogene Technology Inc. All the tissues were subjected to three biological replicates.

### RNA sequencing data analysis

Pair-end RNA-seq reads were first assessed for quality by FastQC v0.10.1 (Andrews, 2010). Trimmomatic was used to remove sequence adapters and reads of low quality (Phred Q < 20) (Bolger *et al*., 2014). High-quality and clean RNA-seq reads were mapped to the reference genome of *Papaver somniferum* HN1 (Yang *et al*., 2021) using bowtie2/2.3.5 (Langmead and Salzberg, 2012). Mapped reads were filtered using Samtools to retain only those that had a mapping quality score of 10 or higher (Samtools ‘view’ command with option ‘-q 10’ to set mapping quality cutoff) (Li *et al*., 2009). Filtered reads were used to construct transcriptome by Cufflinks/2.2.1 (Trapnell *et al*., 2012). Corrplot v0.84 was used to visualize the correlation of gene expression between the different tissues based on scaled normalized read count (TPM) (Wei *et al*., 2017). Subsequently, dimension reduction (PCA analysis) was performed using FactoMineR (Lê *et al*., 2008). After filtering out non-expressed genes (38659), the scaled normalized gene expression matrix was used to perform clustering analysis using WGCNA package (parameters: soft threshold = 14, minModuleSize = 100, MEDissThres = 0.25) (Langfelder and Horvath, 2008).

### ATAC sequencing data analysis

The cleaned reads were mapped to the *Papaver somniferum* genome using BWA (Li and Durbin, 2009) software with ‘mem’ parameters (Yang *et al*., 2021). Mapped reads in sam format were converted to bam format and sorted using Samtools v1.9 and Sambamba ‘markdup’ command was then used to remove PCR duplicates (Li *et al*., 2009, Tarasov *et al*., 2015). Reads with higher mapping quality scores (MAPQ ≥ 10) were employed to perform downstream analysis.

ATAC-seq peak calling was conducted using Genrich with default recommended parameters (v0.6, available at https://github.com/jsh58/Genrich). Intervene was used to intersect different samples to obtain tissue-specific peaks and overlapped peaks (Khan and Mathelier, 2017). The number of reads of each genome region was counted using the ‘multiBamSummary’ script in deepTools v2.0 and correlation and PCA were performed with corrplot and FactoMineR (Lê *et al*., 2008, Ramírez *et al*., 2016, Wei *et al*., 2017).

For each ATAC-seq data set, the peaks were assigned to genes using the R/Bioconductor package ChIPseeker (Yu *et al*., 2015). This program assigns each peak to the closest TSS, whether promoter, downstream, distal intergenic, intron, exon, 5’ UTR, or 3’ UTR, and reports the distance from the peak center to the TSS based on the genome annotations (Yang *et al*., 2021). The TF motif enrichment analysis on ATAC-seq data was performed using the ‘findMotifsGenome.pl’ function of HOMER package (Heinz *et al*., 2010). A potential regulatory TF in opium poppy was selected with expression levels (TPM ≥ 5), uniport annotation (based on *Arabidopsis thaliana* (Mouse-ear cress) [3702]), iTAK annotation, BLASTp results (parameters: -outfmt 6 -best_hit_score_edge 0.05 -best_hit_overhang 0.25 -qcov_hsp_perc 60.0 -max_target_seqs 1, identity ≥ 40, and blast score ≥ 100), and gene module information detected by WGCNA (Langfelder and Horvath, 2008, Madden, 2013, Zheng *et al*., 2016, Consortium, 2019). TF binding sites were performed using the ‘annotatePeaks.pl’ function of HOMER package (Heinz *et al*., 2010). The functional annotation of each transcription factor was performed by UniprotR (Soudy *et al*., 2020).

### Enrichment analysis and Visualization

GO and KEGG enrichment analysis of opium poppy gene was conducted using TBtools (Chen *et al*., 2020, Yang *et al*., 2021).

The filtered, sorted and scaled bam files were converted to the bigwig format for visualization using the BAMscale with default parameters (Pongor *et al*., 2020). Genome browser images were created using the Integrative Genomics Viewer (IGV) v2.8.10 (Thorvaldsdóttir *et al*., 2013) and bigwig files were processed as described above.

Motif binding regions were visualized using ggmsa (v0.0.6, available at https://cran.r-project.org/web/packages/ggmsa/index.html). A Venn diagram for ATAC-seq and RNA-seq samples was generated using R package Venn (v1.9, available at https://cran.r-project.org/web/packages/venn/). The distribution of ATAC-seq peaks was visualized with Circos (Krzywinski *et al*., 2009). The ‘sankeyNetwork’ function in networkD3 (v0.4, available at https://christophergandrud.github.io/networkD3/) was used to create a Sankey diagram. A Heatmap was generated using pheatmap package and dot, bar, pie, and, line plots were all created using ggplot2 package (Wickham, 2016, Kolde, 2019).

### Data availability

ATAC-seq and RNA-seq data generated in this study were deposited in NCBI Sequence Read Archive (SRA) with BioProject accession number PRJNA746779.

## Acknowledgements

This work was supported by the National Natural Science Foundation of China (32125009, 32070663), Chinese Postdoctoral Research Foundation (Grant No. 2020M683514) and the Key Construction Program of the National ‘985’ Project.

